# In silico characterization and phylogenetic analysis of Elaeocarpus ganitrus based on ITS2 barcode sequence

**DOI:** 10.1101/2023.07.15.549003

**Authors:** Jyotsana Kushwaha, Alpana Joshi

## Abstract

Plant molecular systematics relies on using DNA barcodes for studying the evolutionary relationship between species Sequences of the nuclear internal transcribed spacer (ITS) regions have been used widely in molecular phylogenetic studies because of their high variability compared to plastid sequences. Elaeocarpus is a diverse genus within the family Elaeocarpaceae and is widely distributed worldwide among tropical and subtropical climatic zones. Elaeocarpus ganitrus has important medicinal and religious values in India. However, Elaeocarpus ganitrus evolutionary relationship with other Elaeocarpus species is not much explored, especially at the molecular and phylogenetic levels. The present research successfully amplified the nuclear gene ITS2, sequenced and submitted it to NCBI Genbank after using Basic Local Alignment Search Tool (BLAST). Automatic Barcode Gap Discovery (ABGD) and Assemble Species by Automatic Partitioning (ASAP) resulted in different numbers of molecular operational taxonomic units (MOTUs). The lowest score of ASAP (4.5) segregated the sequences into 31 MOTUs with the Threshold dist. value of 0.003524. This study establishes an evolutionary relationship between Elaeocarpus ganitrus and other species belonging to the same genus through the neighbor-joining method. The 38 Elaeocarpus samples were clustered into seven major groups based on ITS2 sequence: Group I is represented by Elaeocarpus ganitrus along with Elaeocarpus sylvestris, Elaeocarpus glabripetalus, Elaeocarpus duclouxii, Elaeocarpus decipiens, and Elaeocarpus zollingeri. Group II is characterized by Elaeocarpus austroyunnanensis and Elaeocarpus glaber. Group III comprises Elaeocarpus sphaericus, Elaeocarpus angustifolius, Elaeocarpus grandis, Elaeocarpus ptilanthus, and Elaeocarpus sphaerocarpus. Three accessions of Elaeocarpus hookerianus are placed in group IV. Elaeocarpus largiflorens and Elaeocarpus thelmae represent group V. Groupr VI contains three species: Elaeocarpus sylvestris, Elaeocarpus dubius, and Elaeocarpus johnsonii. Group VII comprises five species which include Elaeocarpus glabripetalus, Elaeocarpus rugosus, Elaeocarpus tuberculatus, Elaeocarpus hainanensis, and Elaeocarpus angustifolius. The study concludes with the possibility of correctly using the ITS2 gene to identify, discriminate, and documentation of Elaeocarpus ganitrus and other species of the same genus.

## 1. Introduction

*Elaeocarpus* is the most species-rich genus in the family *Elaeocarpaceae* comprising 350 – 400 species ranging from lowland to montane areas of Madagascar, Asia, Australia, and the Pacific islands [1–2]. Most *Elaeocarpus* species are evergreen trees or shrubs, although a few species can occur as epiphytes or lianes, and some are briefly deciduous. *Elaeocarpus ganitrus* is a predominant species of the genus *Elaeocarpus*. It is a large evergreen tree with broad leaves, mostly in subtropical and tropical areas. *Elaeocarpus* tree is famous for its spiritual or aesthetic values and is known as Rudraksha in India. Several phylogenetic studies have been undertaken on the family Elaeocarpaceae and the genus Elaeocarpus, but these were limited in terms of the number of taxon and loci sampling. Phylogenetic relationships within the genus *Elaeocarpus* have been investigated using phylogenetic analysis of nuclear and plastid DNA sequences [3–5]. The nuclear genome is the largest as compared to mitochondrial and plastid genomes. A frequently used part of the nuclear genome for phylogenetic analysis is nuclear ribosomal DNA (nrDNA). Nuclear ribosomal DNA comprises three coding regions (18S, 5.8S, and 26S rDNA) and the non-coding spacer regions (Intergenic spacer (IGS), External transcribed spacer (ETS), and Internal transcribed spacer (ITS) [6]. ITS sequence data for 29 *Elaeocarpus* taxa were analyzed, and the relationships among the genera using nuclear ITS and plastid trnL-trnF sequences from representatives of all accepted genera of *Elaeocarpaceae* were established [3– 4]. Moreover, phylogenetic studies were conducted based on trnL-trnF, trnV-ndhC intergenic spacer, and ITS sequence data of 59 *Elaeocarpus* taxa [5]. Together these molecular studies resolved with some confidence the phylogenetic relationships between most *Elaeocarpaceae* genera. While many clades are robustly resolved, relationships among most *Elaeocarpus* species are unclear due to low sequence divergence.

Initially, DNA barcoding was proposed to assign an unambiguous tag to each species, giving taxonomists a standard method for identifying specimens. DNA barcoding is used to identify species using short-standardized sequences and only requires a small tissue sample [7]. It has recently become an important taxonomic tool because of its precision, reproducibility, and rapidity [8]. Consortium for the Barcode of Life (CBOL) suggested rbcL and matK regions as a standard two-locus barcodes for global plant databases because of their species discrimination ability after comparing the performance of seven candidate barcoding regions, namely atpF-atpH, matK, rbcL, rpoB, rpoC1, psbK-psbI and trnH-psbA [9]. Significant progress has been achieved in the DNA barcoding of higher plants. Plastid DNA barcodes (rbcL, ndhJ, matK, trnL, and trnH-psbA regions) and nuclear DNA barcodes (ITS and ITS2 regions) are commonly used in DNA barcoding of plants [8,10–12]. ITS gene sequence belongs to ribosomal DNA in the nuclear genome and is widely distributed in photosynthetic eukaryotic organisms (except ferns). A large amount of sequence data of the ITS gene has been deposited in GenBank and has become the most common sequence for the phylogeny construction of various crops [13–16]. However, DNA barcodes have been scarcely studied in phylogeny and species identification of plants belonging to the genus *Elaeocarpus*. The aims of the current study were; amplification and sequencing of the ITS region of *Elaeocarpus ganitrus* to study functional annotation and homology modeling of ITS sequence using the Basic Local Alignment Search Tool (BLAST) and Phylogenetic analysis.

## 2. Materials and methods

### 2.1 Plant sampling

The plant samples of *Elaeocarpus ganitrus* have collected in 2023 from Shobhit Institute of Engineering & Technology (SIET), (Deemed-to-be University), Modipuram, Meerut, with the coordinates of the sites, Latitude 29.071274° and Longitude 77.711929° **(Figure 1)**.

**Figure 1.**
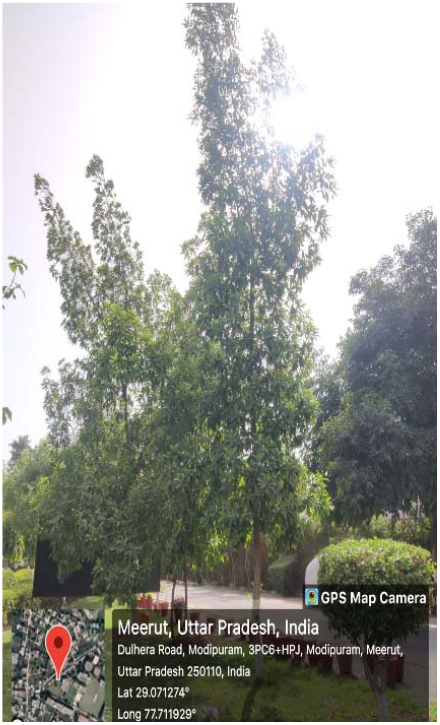
The leaf sample was collected from *Elaeocarpus ganitrus* plant grown at the SIET Campus, Meerut, UP, India.

### 2.2. DNA Extraction, amplification, and sequencing

Young leaf samples were collected and crushed in liquid nitrogen to get fine powder using a sterile mortar and pestle. Total genomic DNA was extracted from about 100 mg of leaf material using a modified CTAB method [17–18]. The extracted DNA quality was further estimated using NanoDrop1000 (Thermo Scientific). The quality of the extracted DNA was also evaluated using 0.8% agarose gel stained with ethidium bromide (1 μg/ml). The specific primer pair were used for the amplification of ITS region (ITS2-F: 5’-ATGCGATACTTGGTGTGAAT-3’ and ITS2-R: 5’-GACGCTTCTCCAGACTACAAT-3’). A PCR reaction mixture of 50 µL comprised the following: template 30–60 ng of DNA, 5 µL PCR buffer (10X), 5 µL of both forward and reverse primers (10 pmol), 5 µL dNTPs (1mM), and 0.35 µL Taq polymerase, and the volume was adjusted with deionized distilled water. PCR-based amplification of ITS barcoding regions was performed in a thermal cycler (Applied Biosystems, USA). The PCR thermal profile was 94 °C for 4 min, followed by 30 cycles of 30 s at 94 °C, 40 s at 55 °C, and 1 min at 72 °C, with a final step of 10 min at 72 °C. The PCR products were examined via electrophoresis in a 1.2% agarose gel containing ethidium bromide and were visualized using an ultraviolet transilluminator. Sequencing was done following the kit’s protocol (BigDye Terminator v3.1, Applied Biosystems). The same PCR primers have been used for sequencing. The reaction has done in ABI 3130xl DNA Analyzers (Applied Biosystems, USA). Each generated consensus sequence of the forward and reverse sequences was submitted to the Basic Local Alignment Search Tool (BLAST) of the National Centre for Biotechnology Information (NCBI) for a homology search (https://blast.ncbi.nlm.nih.gov/Blast.cgi; accessed on 24 June 2023). Sequences were assembled and edited manually using bioedit v7.05. The GenBank accession number for the generated barcode sequences were obtained after the sequences were submitted to the submission portal of NCBI for ITS2 (https://submit.ncbi.nlm.nih.gov/about/genbank/ accessed on 24 June 2023). The final sequences have then been deposited in NCBI GenBank [19].

### 2.3. Sequence alignment and data analysis

The sequence alignment was initially performed by using MUSCLE program of MEGA11 with the default alignment parameters for multiple sequences alignment parameters [20]. In the pairwise distances analyses, the positions containing gaps and missing were eliminated from the data set (complete deletion option). Phylogenetic trees constructed with the Neighbor-joining (N.J.) method according to Kimura 2-Parameter (K2P) model was assessed by MEGA 11 [21]. Evolutionary divergence for each data set and pattern of nucleotide substitution was performed by MEGA 11. For phylogenetic analysis, we used the Neighbor-Joining tree method with 10,000 bootstraps. The number of INDELs for each dataset was identified by deletion/insertion polymorphisms (DIP) analysis in DnaSP v5 [22]. The polymorphic site, genetic diversity indices, and neutrality tests [Fu & Li’s D, Fu & Li’s F, and Tajima’s D were performed using the DnaSP v5 (URL: http://www.ub.edu/dnasp/index_v5.html) [8].

### 2.4. Species delimitation

DNA-based species delimitation was performed using the Automatic Barcode Gap Discovery (ABGD) [23] and Assemble Species by Automatic Partitioning (ASAP) [24]. Automatic Barcode Gap Discovery (ABGD) was conducted using ABGD webserver (URL: https://bioinfo.mnhn.fr/abi/public/abgd/abgdweb.html accessed on June 19, 2023) with default parameters using Kimura 2-Parameter (K2P) model [23]. Assemble Species by Automatic Partitioning (ASAP) was conducted using the ASAP webserver (URL: https://bioinfo.mnhn.fr/abi/public/asap/ accessed on June 23, 2023) using the distance matrix generated through MEGAv11.

## 3. Results and Discussion

### 3.1. Amplification, Sequencing, Multiple Sequence Alignment, and Species Identification

High-quality genomic DNA was isolated from *Elaeocarpus ganitrus* leaf samples, and the 260/280 nm ratio was 1.8 (**Figure 2A**). ITS barcode was amplified from the genomic DNA of four random leaf samples of *Elaeocarpus ganitrus* using gene-specific primer and produced amplicons of around 500 bp (**Figure 2B)**. ITS2 sequence was 95.44 % similar to the sequences deposited in the public database (https://www.ncbi.nlm.nih.gov/nuccore/OR059254.1). The top forty BLASTn scores for the species identification of all *Elaeocarpus* species are presented in **Table 1**.

**Table 1.** The percentage of similarity of *Elaeocarpus ganitrus* with the closest species in the GenBank based on the ITS2 gene sequence.

| Scientific Name | Max Score | Query Cover | E value | Per. ident | Acc. Len | Accession |
| --- | --- | --- | --- | --- | --- | --- |
| *Elaeocarpus sylvestris* | 725 | 95% | 0 | 95.44 | 477 | OR199431.1 |
| *Elaeocarpus austroyunnanensis* | 664 | 86% | 0 | 95.9 | 426 | MW044367.1 |
| *Elaeocarpus braceanus* | 638 | 86% | 0 | 94.72 | 428 | MW044369.1 |
| *Elaeocarpus rugosus* | 604 | 86% | 1.00E-173 | 93.49 | 422 | MW044372.1 |
| *Elaeocarpus dubius* | 564 | 86% | 2.00E-161 | 91.63 | 426 | MW044370.1 |
| *Elaeocarpus dubius* | 545 | 82% | 8.00E-156 | 91.77 | 392 | MW044371.1 |
| *Elaeocarpus sylvestris* | 549 | 81% | 6.00E-157 | 92.19 | 390 | MW044374.1 |
| *Elaeocarpus tuberculatus* | 501 | 76% | 2.00E-142 | 91.87 | 693 | KP748174.1 |
| *Elaeocarpus variabilis* | 549 | 76% | 6.00E-157 | 94.02 | 624 | KP748173.1 |
| *Elaeocarpus duclouxii* | 542 | 71% | 1.00E-154 | 95.36 | 357 | KP096061.1 |
| *Elaeocarpus decipiens* | 534 | 67% | 2.00E-152 | 96.6 | 320 | KP096060.1 |
| *Elaeocarpus sphaerocarpus* | 405 | 66% | 1.00E-113 | 90.25 | 399 | KR532067.1 |
| *Elaeocarpus glaber* | 486 | 65% | 5.00E-138 | 94.65 | 511 | KJ675654.1 |
| *Elaeocarpus stipularis* | 475 | 65% | 1.00E-134 | 94.04 | 516 | KJ675666.1 |
| *Elaeocarpus hylobroma* | 435 | 65% | 2.00E-122 | 92.11 | 613 | KJ675680.1 |
| *Elaeocarpus ruminatus* | 435 | 65% | 2.00E-122 | 92.11 | 615 | KJ675662.1 |
| *Elaeocarpus hookerianus* | 424 | 65% | 4.00E-119 | 91.48 | 624 | KJ675656.1 |
| *Elaeocarpus grandis* | 420 | 65% | 5.00E-118 | 91.17 | 616 | KJ675655.1 |
| *Elaeocarpus largiflorens* | 412 | 65% | 9.00E-116 | 90.88 | 602 | KJ675681.1 |
| *Elaeocarpus thelmae* | 412 | 65% | 9.00E-116 | 90.85 | 517 | KJ675670.1 |
| *Elaeocarpus angustifolius* | 412 | 65% | 9.00E-116 | 90.85 | 580 | KJ675645.1 |
| *Elaeocarpus pulchellus* | 411 | 65% | 3.00E-115 | 90.85 | 610 | KJ675684.1 |
| *Elaeocarpus ptilanthus* | 409 | 65% | 1.00E-114 | 90.54 | 617 | KJ675683.1 |
| *Elaeocarpus sphaericus* | 409 | 65% | 1.00E-114 | 90.54 | 617 | KJ675679.1 |
| *Elaeocarpus johnsonii* | 409 | 65% | 1.00E-114 | 90.54 | 617 | KJ675657.1 |
| *Elaeocarpus sphaericus* | 409 | 65% | 1.00E-114 | 90.54 | 616 | KJ675647.1 |
| *Elaeocarpus largiflorens* | 407 | 65% | 4.00E-114 | 90.57 | 612 | KJ675658.1 |
| *Elaeocarpus bifidus* | 399 | 65% | 7.00E-112 | 90.25 | 612 | KJ675650.1 |
| *Elaeocarpus dongnaiensis* | 490 | 65% | 4.00E-139 | 94.97 | 493 | KJ675653.1 |
| *Elaeocarpus sylvestris* | 488 | 65% | 1.00E-138 | 94.95 | 486 | KJ675667.1 |
| *Elaeocarpus zollingeri* | 488 | 65% | 1.00E-138 | 94.95 | 380 | LC600913.1 |
| *Elaeocarpus glabripetalus* | 483 | 65% | 7.00E-137 | 94.64 | 598 | KF186469.1 |
| *Elaeocarpus hainanensis* | 425 | 65% | 1.00E-119 | 91.46 | 632 | KF186468.1 |
| *Elaeocarpus rugosus* | 422 | 65% | 1.00E-118 | 91.22 | 401 | KR532058.1 |
| *Elaeocarpus angustifolius* | 407 | 65% | 4.00E-114 | 90.51 | 619 | KJ675646.1 |
| *Elaeocarpus sylvestris* | 473 | 64% | 4.00E-134 | 94.81 | 608 | KP093064.1 |
| *Elaeocarpus hookerianus* | 407 | 64% | 4.00E-114 | 91.23 | 518 | DQ448688.1 |
| *Elaeocarpus angustifolius* | 405 | 64% | 1.00E-113 | 90.91 | 526 | DQ448689.1 |
| *Elaeocarpus largiflorens* | 396 | 64% | 9.00E-111 | 90.61 | 595 | DQ448686.1 |
| *Elaeocarpus williamsianus* | 394 | 64% | 3.00E-110 | 90.58 | 616 | DQ448691.1 |

**Figure 2.**
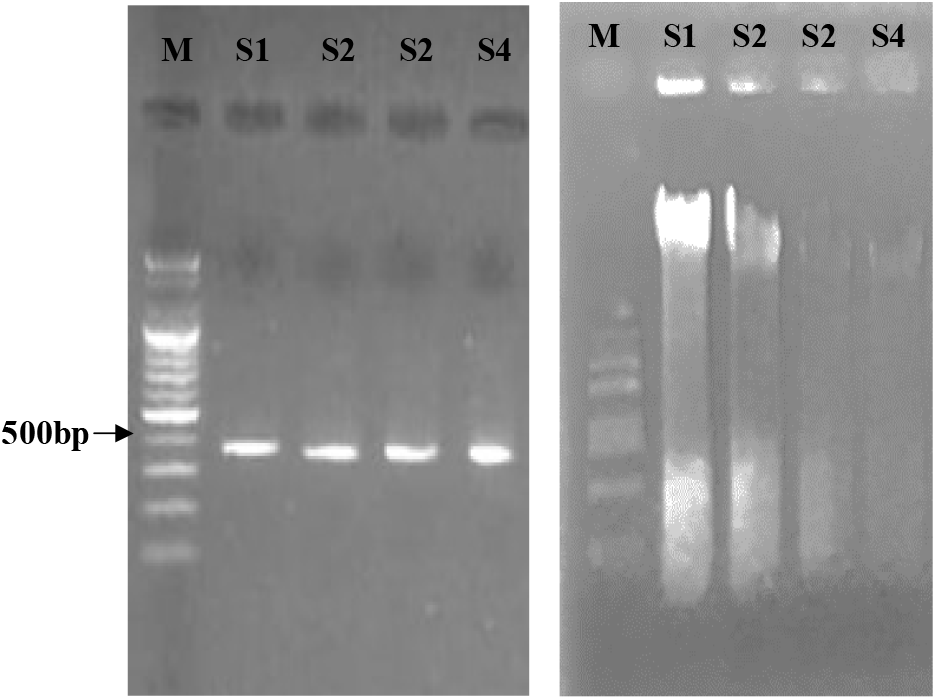
(A) Agarose gel electrophoresis (0.8 %) showing bands of genomic DNA isolated from leaf samples of *Elaeocarpus ganitrus*. (B) Agarose gel electrophoresis (1.2%) showing PCR amplified band of the ITS2 barcoding region (∼ 500 bp). M: 1 kb plus DNA ladder, S1, S2, S3, and S4: leaf samples from *Elaeocarpus ganitrus*.

### 3.2. DNA sequences analysis

This study retrieved 58 ITS2 sequences of genus *Elaeocarpus* from the NCBI Nucleotide database (https://www.ncbi.nlm.nih.gov/) for further analysis. All retrieved sequences were evaluated critically, and any low-quality sequences were removed. Criteria used to filter the sequences containing <3% ambiguous base ‘N’ [25]. After blasting and editing in MEGA v11, the consensus length of ITS2 was found to be 328 bp. This study evaluates the substitution of different bases in analyzed regions on entire codon positions (1st+ 2nd + 3rd nucleotide). Substitution patterns and rates were estimated under the Tamura model [26]. The substitution rate from T to C and A to G was noted in ITS2 region was 15.49 % **(Table 2)**.

**Table 2.** The maximum likelihood estimate of substitution matrix in the constructed phylogenetic tree of genus *Elaeocarpus sp*. ITS2 gene sequence.

|  | A | T/U | C | G |
| --- | --- | --- | --- | --- |
| A | - | 4.75 | 4.75 | **15.49** |
| T/U | 4.75 | - | **15.49** | 4.75 |
| C | 4.75 | **15.49** | - | 4.75 |
| G | **15.49** | 4.75 | 4.75 | - |

**NOTE-** Each entry is the probability of substitution (*r*) from one base (row) to another base (column). Substitution patterns and rates were estimated under the Tamura (1992) model [26]. Rates of different transitional substitutions are shown in **bold**, and those of transversional substitutions are shown in *italics*. Evolutionary analyses were conducted in MEGA11.

### 3.3. Genetic diversity

The distribution of the four bases and mean nucleotide base frequencies observed for the ITS2 sequence were showed in **Table 3**. Polymorphism site analysis of the ITS2 barcode sequence was conducted using DnaSP Version 6.12. The ITS2 sequences had 43 mutation sites and 36 segregating sites. The basic indicators of genetic diversity were calculated in accordance with pairwise nucleotide differences and nucleotide diversity. The sequence analysis exhibited 105 monomorphic sites and 36 polymorphic sites. Neutrality tests verified the significance of genetic diversity; Tajima’s D, Fu & Li’s D, and Fu & Li’s F test statistics. Tajima’s D, Fu’s F.S., and Fu & Li’s F test statistics were statistically negative but not significant (PD>D0.10). The results showed that these populations were stable, with no recent bottleneck or rapid population expansion. However, the Fu and Li’s D test value was negative and statistically significant (PD<D0.10), which showed that these populations had experienced a recent population expansion [27]. To observe nucleotide mismatch distribution among different sequences of *Elaeocarpus* species, DNA sequences were analyzed for population size changes, enriching the results of genetic diversity among species. All results showed significant genetic variation in *Elaeocarpus* species for the ITS2 sequence **(Figure 3A, Table 4)**. The automated clustering analyses were carried out with 58 individual sequences of *Elaeocarpus* species using the Automatic Barcode Gap Discovery (ABGD) method. ABDG method could not detect significant barcoding gaps due to the overlapping of intra- and interspecific distances **(Figure 3B)**. The same data is represented as ranked ordered values **(Figure 3C)**. The pairwise interspecific distances in the ITS barcodes ranged from 0.0069 to 0.0788 **(Figure 3D)**.

**Table 3.** The nucleotide compositional analysis of candidate nucleotide sequences in *Elaeocarpus* species.

| Sequences | Base content |  |  |  |  |  |  |  |  |  |  |  |
| --- | --- | --- | --- | --- | --- | --- | --- | --- | --- | --- | --- | --- |
|  | T | C | A | G | GC | GC-1 | GC-2 | GC-3 | AT | AT-1 | AT-2 | AT-3 |
| ITS2 | 15.91 | 37.74 | 15.93 | 30.39 | 68.13 | 67.22 | 70.129 | 67.09 | 31.84 | 32.76 | 29.87 | 32.90 |

**Table 4.** The analysis of variation of ITS sequences in *Elaeocarpus* plants.

| DNA Sequence Polymorphism |  | Neutrality Test |  |
| --- | --- | --- | --- |
| Number of sequences | 58 | Tajima's D | -1.16266 |
| Number of polymorphic (segregating) sites (S) | 36 | Fu and Li's D test statistic | -0.93964* |
| Total number of mutations, Eta | 43 | Fu and Li's F test statistic | -1.21841 |
| Haplotype (gene) diversity, Hd | 0.961 | Fu's Fs statistic | -12.726 |
| Nucleotide diversity, Pi | 0.04454 |  |  |
| Theta (per site) from Eta | 0.06588 |  |  |
| Sites with alignment gaps or missing data | 187 |  |  |
| Invariable (monomorphic) sites | 105 |  |  |
| Variable (polymorphic) sites | 36 |  |  |
| Singleton variable sites | 8 |  |  |
| Parsimony informative sites | 28 |  |  |

**Figure 3.**
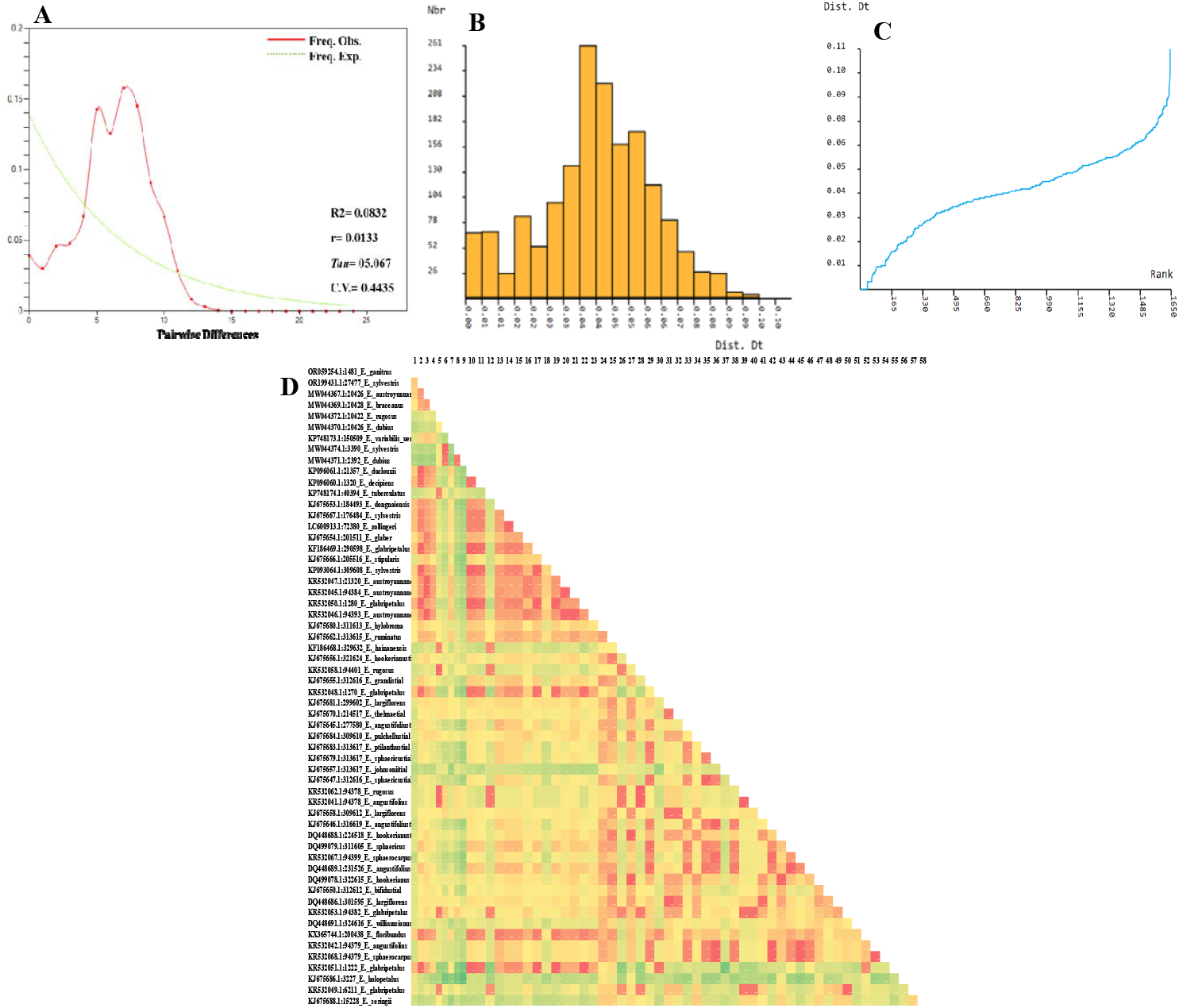
Genetic diversity of ITS2 barcode sequences of genus *Elaeocarpus*. (A) Frequencies of the observed and expected pairwise differences (the mismatch distribution) based on ITS2 sequence of *Elaeocarpus ganitrus*. The X-axis shows the pairwise differences, and the Y-axis shows the frequency. R2 Ramos-Onsins and Rozas statistics, r Raggedness statistic, Tau Date of the Growth or Decline measured of mutational time, C.V. Coefficient of variation. (B) Histogram showing pairwise distance divergence (%) generated by Automatic Barcode Gap Discovery (ABGD) after the input of distance data belonging to 58 ITS2 sequences of the *Elaeocarpus ganitrus*. Nbgroups is the number of species as identified by ASAP in the corresponding partition (C) Ranked distances and (D) Heat map for the pairwise genetic distance between *Elaeocarpus* populations.

This sequence set was partitioned into subsets independently by ASAP [24]. ASAP is based on an algorithm using only pairwise genetic distances to reduce the computational time for phylogenetic reconstruction. For each partition, the number of molecular operational taxonomic units (MOTUs) is identified, and W-values determine the ASAP score and the ‘threshold distance, which is the limit value of genetic divergence for which two sequences are considered to belong to different MOTUs **(Table 5)**. The lowest score of ASAP (4.5) is considered best, and segregated the sequences into 31 MOTUs when the Threshold dist. value was 0.003524. The sequence partition with the ASAP score (5) distributed the sequences into 32 and 30 MOTUs with the Threshold dist. values 0.003423 and 0.003656, respectively. The sequence partition with the ASAP score (6.5) partitioned the sequences into 24 MOTUs with the Threshold dist. value 0.007757. The sequence partition with the ASAP score (7) distributed the sequences into 33 MOTUs and the Threshold dist. value was 0.003345, whereas sequence partition with the ASAP score (8) distributed the sequences into 35 and 34 MOTUs with the Threshold dist. values 0.001648 and 0.003301, respectively. The sequence partition with the ASAP score (9) distributed the sequences into 27 and 26 MOTUs with the Threshold dist. values 0.006857 and 0.006978, respectively. The sequence partition with the highest ASAP score (10.5) distributed the sequences into 29 MOTUs and the Threshold dist. Value was 00.005153 **(Figure 4, Table 5)**.

**Table 5.**
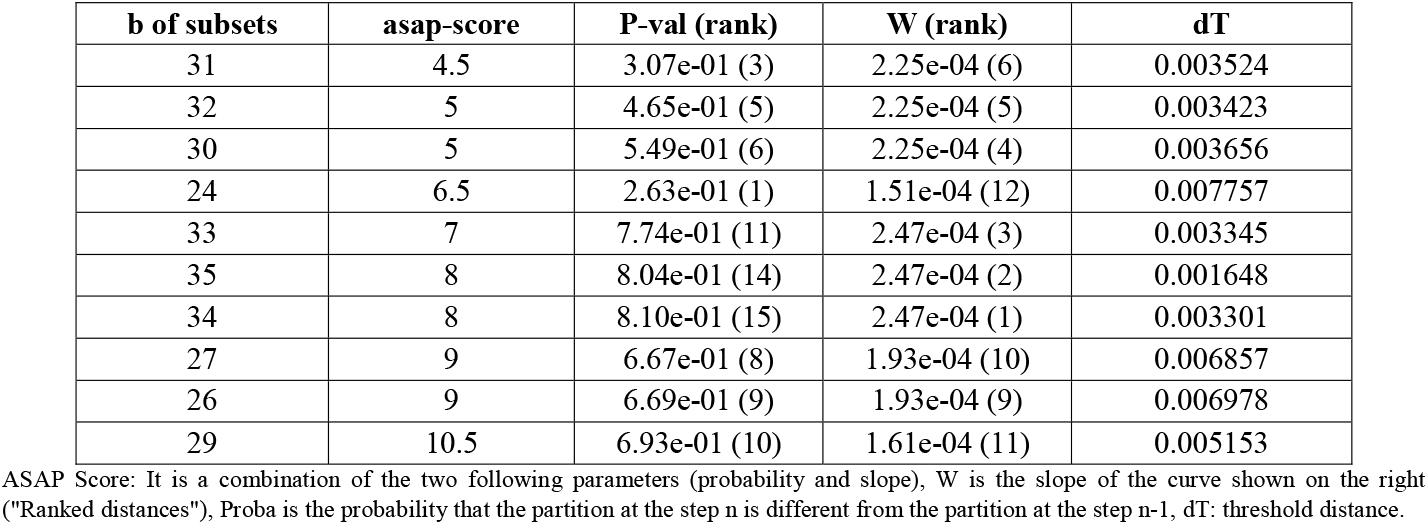
ASAP identifies different partitions based on ITS sequences in *Elaeocarpus* plants.

| b of subsets | asap-score | P-val (rank) | W (rank) | dT |
| --- | --- | --- | --- | --- |
| 31 | 4.5 | 3.07e-01 (3) | 2.25e-04 (6) | 0.003524 |
| 32 | 5 | 4.65e-01 (5) | 2.25e-04 (5) | 0.003423 |
| 30 | 5 | 5.49e-01 (6) | 2.25e-04 (4) | 0.003656 |
| 24 | 6.5 | 2.63e-01 (1) | 1.51e-04 (12) | 0.007757 |
| 33 | 7 | 7.74e-01 (11) | 2.47e-04 (3) | 0.003345 |
| 35 | 8 | 8.04e-01 (14) | 2.47e-04 (2) | 0.001648 |
| 34 | 8 | 8.10e-01 (15) | 2.47e-04 (1) | 0.003301 |
| 27 | 9 | 6.67e-01 (8) | 1.93e-04 (10) | 0.006857 |
| 26 | 9 | 6.69e-01 (9) | 1.93e-04 (9) | 0.006978 |
| 29 | 10.5 | 6.93e-01 (10) | 1.61e-04 (11) | 0.005153 |

ASAP Score: It is a combination of the two following parameters (probability and slope), W is the slope of the curve shown on the right ("Ranked distances"), Proba is the probability that the partition at the step n is different from the partition at the step n-1, dT: threshold distance.

**Figure 4.**
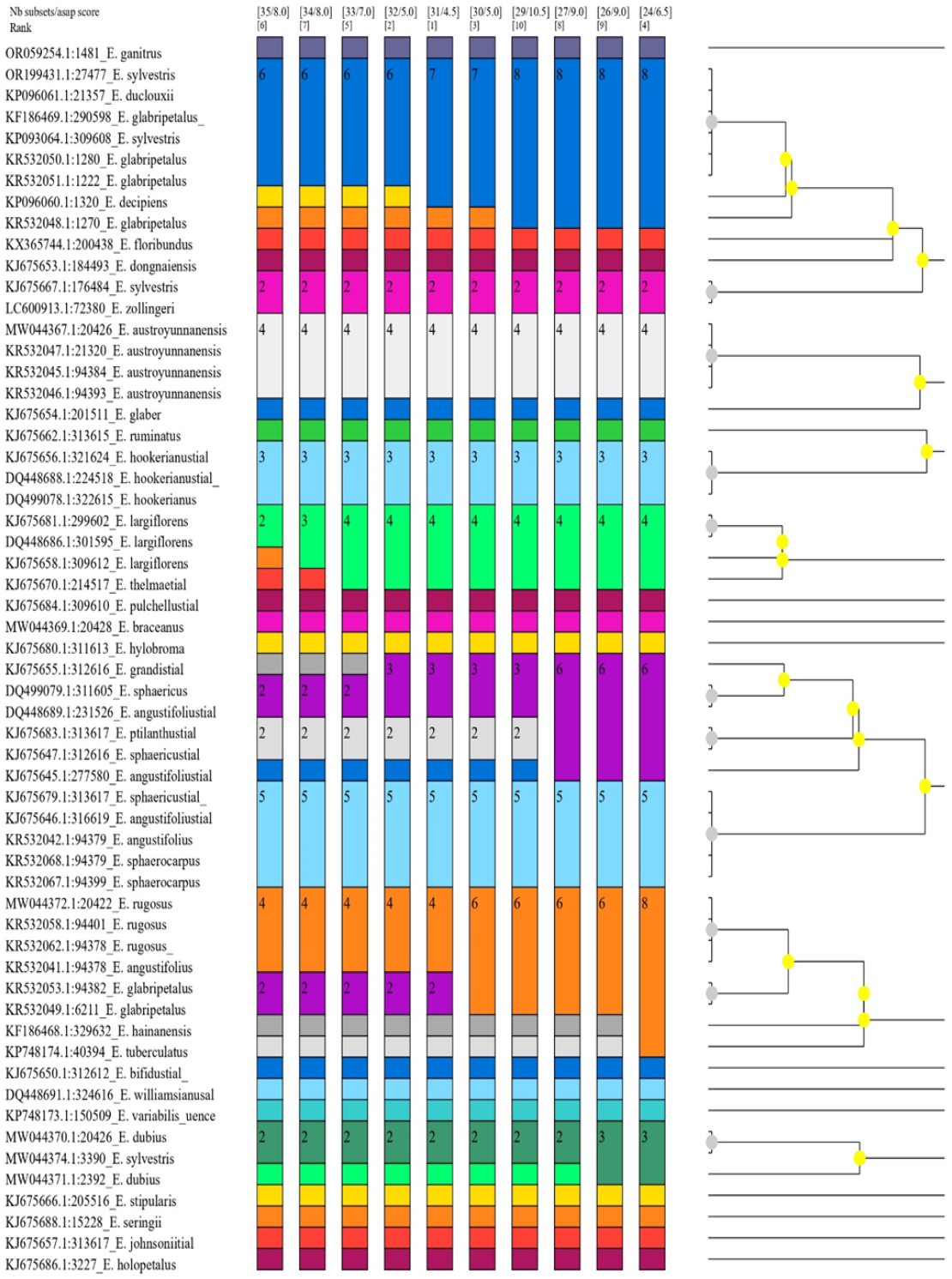
Output results obtained with the ASAP method. Graphical output showing the different delimitations together with the ultrametric clustering tree; each column represents a partition, and the colors represent the molecular operational taxonomic units (MOTUs).

### 3.4. Phylogenetic analysis

This study uses the Neighbor-Joining method and Kimura 2-parameter model to study the evolutionary relationship of *Elaeocarpus ganitrus* within the genus *Elaecarpus* based on ITS2 sequences. A few molecular phylogenetic studies have been conducted on the *Elaeocarpaceae* family [2–5,28]. Within the genus *Elaeocarpus*, seven major clades are identified based on ITS2 sequences of 38 *Elaeocarpus* species (Figure 5). *Elaeocarpus ganitrus* (ORO59254.1) is placed in group I with bootstrap support of 46% and grouped with *Elaeocarpus sylvestris* (OR199431.1, KJ675667.1, and KP093064.1), *Elaeocarpus glabripetalus* (KF186469.1, KR532050.1, KR532048.1, and KR532051.1), *Elaeocarpus duclouxii* (KP096061.1), *Elaeocarpus decipiens* (KP096060.1) and *Elaeocarpus zollingeri* (LC600913.1). All the four accessions of *Elaeocarpus austroyunnanensis* (MW044367.1, KR532047.1, KR532046.1, and KR532045.1) formed a group II along with *Elaeocarpus glaber* (KJ675654.1) and resolved with 57 % bootstrap support. Group III resolved with 85 % bootstrap support and comprises *Elaeocarpus sphaericus* (DQ499079.1, KJ675647.1, and KJ675679.1), *Elaeocarpus angustifolius* (KJ675645.1, KJ675646.1, DQ448689.1, and KR532042.1), *Elaeocarpus grandis* (KJ675655.1), *Elaeocarpus ptilanthus* (KJ675683.1), *Elaeocarpus sphaerocarpus* (KR532067.1 and KR532068.1). All the three accessions of *Elaeocarpus hookerianus* (KJ675656.1, DQ448688.1 and, DQ499078.1) are placed in group IV with 91% bootstrap support. Group V comprises *Elaeocarpus largiflorens* (KJ675681.1, KJ675658.1, and DQ448686.1) and *Elaeocarpus thelmae* (KJ675670.1) resolved with 72 % bootstrap support. One accession of *Elaeocarpus sylvestris* (MW044374.1) is placed in group VI with 40 % bootstrap support along with *Elaeocarpus dubius* (MW044370.1 and MW044371.1), and *Elaeocarpus johnsonii* (KJ675657.1). Group VII comprises five species resolved with 79 % bootstrap support which includes *Elaeocarpus glabripetalus* (KR532053.1 and KR532049.1), *Elaeocarpus rugosus* (MW044372.1, KR532058.1, and KR532062.1), *Elaeocarpus tuberculatus* (KP748174.1), *Elaeocarpus hainanensis* (KF186468.1), and *Elaeocarpus angustifolius* (KR532041.1). The ITS sequence has been used in previous phylogenetic studies of the *Elaeocarpaceae* family [3–4] but has not yet provided a satisfactory resolution of some of the clades. The phylogeny of the family *Elaeocarpaceae* was analyzed based on nuclear ITS sequences of 50 species representing the 12 genera of the *Elaeocarpaceae* family using Parsimony and Bayesian methods [2–5]. The molecular phylogenetic study in *Elaeocarpus* [3] investigated the phylogeny of Australian species based on analyses of nuclear ITS sequences of 32 species of *Elaeocarpus* using Parsimony and Likelihood analyses. Furthermore, the phylogenetic relationships among *Elaeocarpus* species of Australia were conducted with a much-expanded dataset [5]. Afterward, Phoon [2] demonstrated that *Elaeocarpus* is monophyletic, based on 114 species of *Elaeocarpus*. Evolutionary studies concerning *Elaeocarpus* have unlocked new perspectives on the plant’s evolutionary history, which could not be achieved through morphological studies. Based on morphological data, Ganitrus clade formed two main clades; one clade comprises; *Elaeocarpus angustifolius, Elaeocarpus sphaericus, Elaeocarpus ptilanthus* and *Elaeocarpus kaniensis*, and the other contains *Elaeocarpus polydactylus, Elaeocarpus nubigenus, Elaeocarpus murukkai*, and *Elaeocarpus dolichostylus*. Although the phylogenetic relationship between the different species of *Elaeocarpus* has been studied by various researchers using molecular and morphological markers, however studies at the molecular phylogenetic level to understand the evolutionary history of *Elaeocarpus ganitrus* and other species of the genus *Elaeocarpus* are limited. The present research can be a foundation for further investigations into *Elaeocarpus* molecular systematics and their evolutionary history using more diverse phylogenetic markers and species.

**Figure 5.**
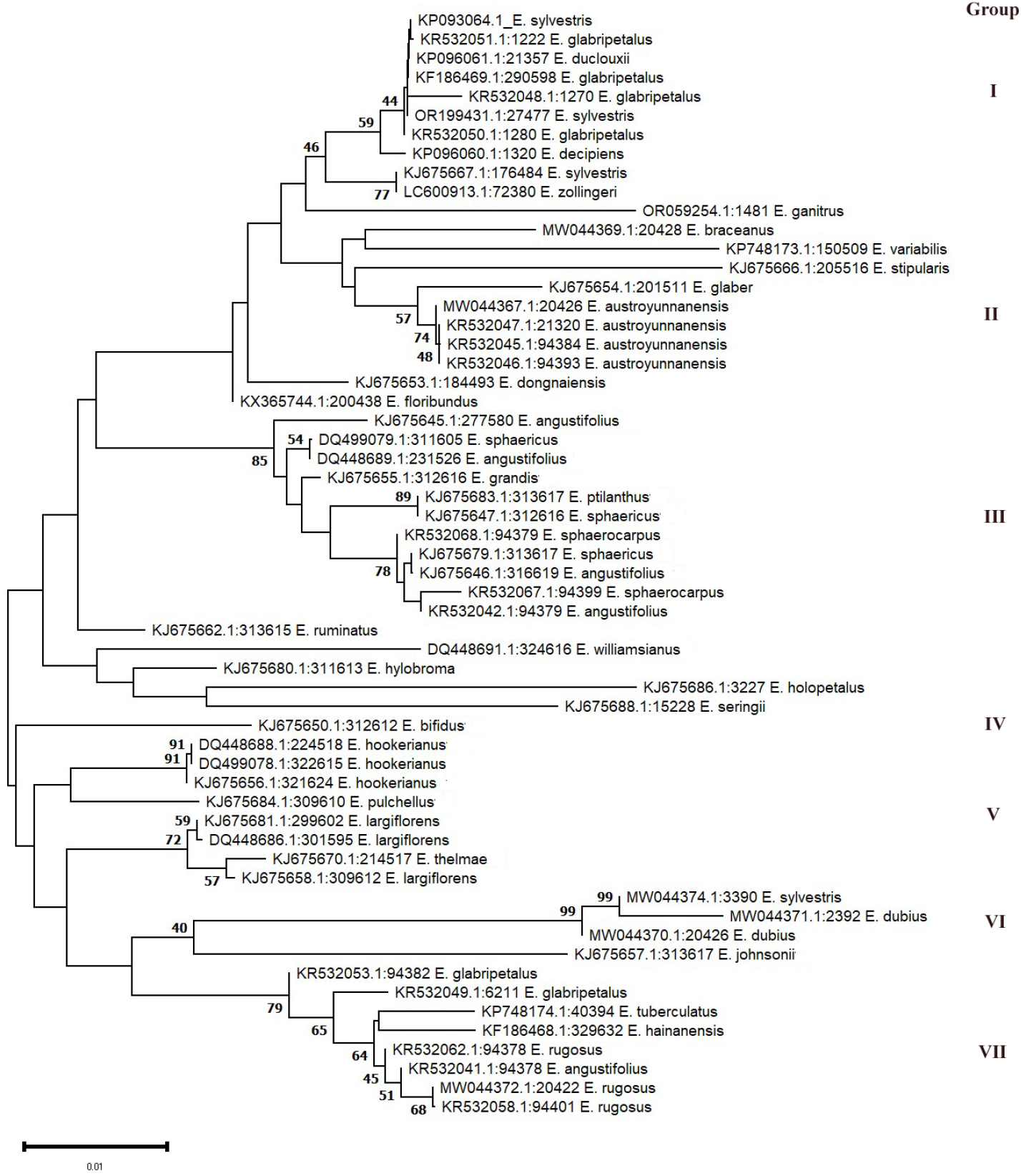
Neighbor-joining tree of *Elaeocarpus* based on *ITS2* sequences using Tamura 3-parameter method. The Numbers on the branches represent more than or equal to 40 percent support after the 10,000 bootstrap replications test. Evolutionary analyses were conducted in MEGA11.

## Conclusion

DNA barcodes can be utilized for species identification by amplifying DNA fragments common to all species. The nuclear DNA comprises several hundred to thousands of tandemly aligned copies of gene cassettes. ITS region has been popular for phylogenetic reconstruction because of its abundant copies and semi-universal primers. The high levels of variation within the non-coding parts of nuclear DNA are advantageous for phylogenetic studies, even in population-level genetic diversity studies. This study is the first attempt to amplify the ITS2 region of *Elaeocarpus ganitrus* and recommends using ITS2 as a barcode for authenticating plants belonging to the genus *Elaeocarpus*. The present study provides a better resolution at the species level and largely agrees with the previously hypothesized phylogenetic relationships of the *Elaeocarpus*. The phylogenetic analyses suggest that the identification ability of ITS2 can be improved in combination with other barcodes. A comprehensive approach and multiple DNA markers could also be employed to understand the relationships between *Elaecarpus ganitrus* and other *Elaecarpus* species.

## Conflicts of Interest

The authors declare no conflict of interest.

## Funding Statement

This research received no specific grant from any funding agency.

## Notes

### Competing Interest Statement

The authors have declared no competing interest.

